# DBP2Vec: Predicting DNA-binding proteins directly using pre-trained protein language model

**DOI:** 10.1101/2022.07.30.502114

**Authors:** Chao Wei, Zhiwei Ye, Panru Wang, Wuyang Lan

## Abstract

DNA-binding proteins (DBPs) play a crucial role in numbers of biological processes and have received wide attention in recent years. Meanwhile, the rapid development of sequencing technologies lead to the explosive growth of new protein sequences, it is highly desired to develop a fast and accurate method for DNA-binding proteins prediction. Experimental methods such as chromatin immunoprecipitation on microarray (ChIP-chip) and X-ray crystallography are highly accurate but expensive and time-consuming. To address this issue, many computational methods have been proposed, they usually exploit multiple information about protein sequence, e.g., sequence composition information, physicochemical properties, evolutionary information, structural information, etc. Despite the effectiveness of these approaches, they heavily depend on prior biological knowledge and undergo a very complex process of feature extraction. In view of these shortcomings, here, we present a novel method, named DBP2Vec, to predict DNA-binding proteins directly from pre-trained protein language model (e.g., ESM-1b) which effectively encode biological properties without any prior knowledge by virtue of deep representation learning (e.g., BERT) on large protein sequences databases (e.g., UniParc). Tests on two DBPs benchmark datasets (e.g., PDB296, UniSwiss-Tst) demonstrate that our proposed method significantly outperforms existing state-of-the-art methods. The source code and the datasets used in the paper are publicly available at: https://github.com/hgcwei/DBP2Vec.

## Introduction

DNA-binding proteins (DBPs) is a kind of proteins that play critical roles in various molecular functions, e.g., DNA replication and repair, the detection of DNA damage, transcriptional control, chromatin stability, etc [1, 2]. Furthermore, many studies demonstrate that DBPs are associated with numerous diseases like cancer [3], neurodegenerative disease [4], etc. Accurately predicting DNA-binding proteins are not only meaningful to theoretical research, but also real applications. Tremendous experimental methods such as chromatin immunoprecipitation on microarrays (ChIP-chip) [5], X-ray crystallography [6], and filter binding assays [7] can accurately identify DBPs, but at the expense of time and money. It is highly desired to develop an automatic computational method for rapidly and accurately predicting DBPs.

In recent years, many computational methods have been proposed to identify DBPs. These methods mainly exploit at least one among the five kinds of information extracted from protein sequence, including 1) sequence composition information, this is a basic feature which is widely used for biological sequence function or structure classification [8, 9], the commonly used features includes Amino Acid Composition (AAC) [10], Pseudo Amino Acid Composition (PseAAC) [11], etc. 2) physicochemical properties, this kind of features can also carry information about the biological function of protein sequence, which is useful to distinguish DBPs and non-DBPs. 3) evolutionary information, this kind of information is based on sequence conservation that protein sequence belonging to the same class often have similar sequence composition, e.g., motifs, transition, etc. The commonly used evolutionary information is position-specific scoring matrix (PSSM). 4) structural information, Secondary structure and 3d structure also plays an important role in the function of DBPs [12], e.g., many DBPs show obvious preference of certain secondary structure motifs, such as helixturn-helix and coil-helix-coil [13].

Chou’s PseAAC has been successfully applied to nearly all areas of computational proteomics [14–16] for its effectiveness to reflect the global sequence order information with comparison to AAC by modeling the interactions between distant residues of protein sequence. Some methods even directly used this method to extract sequence composition information for DBPs prediction. For example, Lin et al. [17] proposed a method, called iDNA-Prot to incorporate the grey model parameters into PseAAC. Cai and Lin [18] formulated a 40-dimensional feature vector from PseAAC to represent DBPs. Rahman et al. [19] present a model called DPP-PseAAC to exploit sequence composition information such as AAC, n-gapped-dipeptides (nGDip), and position-specific n-grams (PSN) for DBPs prediction. Moreover, some methods also used only 3D structure of protein sequence to predict DBPs. The method of Ahmad et al. [20] represents proteins with 62 structural features including the protein’s net charge, electric dipole moment and quadrupole moment tensors. Similarly, Nimrod et al. [21] computed various structural characteristics of proteins from their average surface electrostatic potentials, dipole moments and cluster-based amino acid conservation patterns. Doubtless, fusion of multiple information aforementioned can bring better prediction performance than only using one. Nanni et al. [22] used AAC and quasi residue couple (QRC) to extract sequence composition information, and physicochemical properties were also extracted by the autocovariance approach (AC). Zhang et al. [23] used 14 kinds of physicochemical property, protein secondary structural information, and evolutionary information. Chowdhury et al. [24] proposed a method called iDNAProt-ES which utilizes both evolutionary information and sequence-driven structural information of protein sequence to identify DBPs. Liu et al. [25] present a method called PseDNA-Pro to utilize AAC, PseAAC and physicochemical distance transformation for DBPs prediction. TargetDBP [26] integrated four features including AAC, pseudo predicted relative solvent accessibility (PsePRSA), pseudo position-specific scoring matrix (PsePSSM), and pseudo predicted probabilities of DNA-binding sites (PsePPDBS) for DBPs prediction. Later, they present its updated version TargetDBP+ [27], where a new feature called amino acid one-hot matrix (AAOHM) is added.

Despite the outstanding performance achieved by the aforementioned methods, they still have much room to improve. First, the sequence composition information extracted by PseAAC can only model residue-residue interactions in a short range, which limits their capability to reflect global sequence order information. Second, the feature extraction of these methods is complex and computationally expensive, e.g., the generation of PSSM requires PSI-BLAST [28] search in a huge protein database, the protein secondary structure and relative solvent accessibility information requires extra prediction in advance by using specific software, e.g., SSPro, ACCPro [29]. Third, the fusion of multiple heterogeneous features may bring redundancy and noise which reduce the efficiency of the model [30]. Fourth, for the methods that use the 3D structure information, it is only applicable when the high-resolution 3D structure of protein is available, however, there only exists a small portion of samples that have 3D structure data, which will limit their further application [27]. Very recently, the transformer model [31] transferred to a massive amount of protein sequences using self-supervision approaches have demonstrated a significant potential for harnessing the power of big data and substantially improve the prediction performance on various tasks [32–34]. They showed that without any prior knowledge, the pre-trained models could effectively learn fundamental properties of proteins such as secondary structure, contacts, and biological activity. This gives us an intuition that we can examine its performance on the task of DBPs prediction, where the structural information is required.

Based on the above analysis, we present a simple and accurate DBPs predictor, named DBP2Vec, directly using the pre-trained protein language model (e.g., ESM-1b [33]). To be specific, for each protein sequence, a feature vector is firstly generated directly from ESM-1b, and then the optimal feature set is extracted from the generated feature vector by using a mutual information-based feature selector [35], which is finally fed into a support vector machine (SVM) classifier for DBPs prediction. Compared with existing state-of-the-art methods, DBP2Vec is not only simpler (exploit only embedding from ESM-1b), but also more accurate when test on two benchmark DBPs datasets. To our best knowledge, this is the first attempt to introduce pre-trained protein language model to DBPs prediction.

## Materials and Methods

### Datasets

We adopted two benchmark datasets strictly selected by TargetDBP, and TargetDBP+ to test our proposed method, e.g., PDB used in TargetDBP and UniSwiss in TargetDBP+. For PDB dataset, two subsets, e.g., PDB-Tr and PDB296 are constructed for model training and testing, respectively. To be specific, all DBP chains from PDB [36] (as of May 12, 2018) are firstly extracted and each DBP chain is classified into the positive class in PDB or contains one DNA-binding residue at least. Then, the CD-hit [37] software [29] is used to remove the redundant protein chains such that any two chains have no more than 25% sequence identity. To ensure no fragment in the final dataset, any chain with less than 50 residues in length is removed. Besides, the proteins containing the residue ‘X’ are also removed since they include unknown residue. Finally, a total of 1,200 non-redundant DBPs are obtained and selected as positive samples. Following the same procedure above, 16,058 non-DBPs are obtained after redundancy removal at sequence identity less than 25%. Finally, 1,052 DBPs and 1,052 non-DBPs are selected as training set (PDB-Tr), and 148 DBPs and 148 non-DBPs are selected as independent testing set (PDB296). For UniSwiss dataset, 32,890 DBPs are downloaded firstly in the UniProtKB/Swiss-Prot1 database [38] (up to October 5, 2020) at https://www.uniprot.org/keywords/KW-0238, and then the training set UniSwiss-Tr and independent testing set UniSwiss-Tst are constructed by following the same procedure as PDB. Finally, 4,500 DBPs and 4,500 non-DBPs are included in UniSwiss-Tr and 381 DBPs and 381 non-DBPs are included in UniSwiss-Tst.

### Pre-trained protein language model

Distributed representation has been proved to be a successful data representation approach in natural language processing in the past years. Compared with one-hot encoding, distributed representation contains more semantic information about language context and more suitable for tasks such as sentiment classification [39], text classification [40]. Indeed, biological sequences (e.g., DNA, RNA and protein sequence) show very similar characteristics with natural language, many biologists believe that biological sequences are not merely one-dimensional string of symbols, but encode a lot of useful information about molecular structure and functions in themselves [41]. Hence, it is a natural idea to introduce distributed representation in natural language processing to biological sequence analysis. It is firstly introduced by [42] to protein family classification and a prediction accuracy of 99% is achieved, then it is pervasive in a wide range of applications for biological sequences analysis, e.g., protein secondary structure prediction [43], RNA-protein binding sites prediction [44]. Existing studies showed that the pre-trained models could substantially improve the predictive performance on various tasks [32–34].

In this paper, we introduce the pre-trained protein language model to DBPs prediction for the first time. We apply the pre-trained ESM-1b [33] model for its successful applications to a number of biological sequence analysis, including sequence-level prediction, and residue-level prediction. As shown in Fig 1, the main pipeline of DBP2Vec contains three steps, including feature extraction, feature selection and classification construction. During the process of feature extraction, the dataset (e.g., DBPs, non-DBPs) is split into a training set and testing set, then both of them are fed into pre-trained protein language model, e.g., ESM-1b, to get the embedding feature vector, each sample is converted into a 1280-dimensial feature vector. Then, a feature selection process is performed to eliminate the features that is irrelevant to DBPs prediction. We used univariate feature selection like mutual information for its efficiency. During the process of classification model construction, we adopt the prevalent used libsvm [45] for this goal, the optimal feature vectors of training and testing set are fed into the SVM classifier to train and test the classification model. The radial basis function is selected as the kernel function, and the MCC value is used as the function to optimize the parameters (C and gamma). Here, for all training data, the optimal values of C = 2.0, gamma = 0.5 are obtained by grid search method (run grid.py script in libsvm).

**Figure 1.**
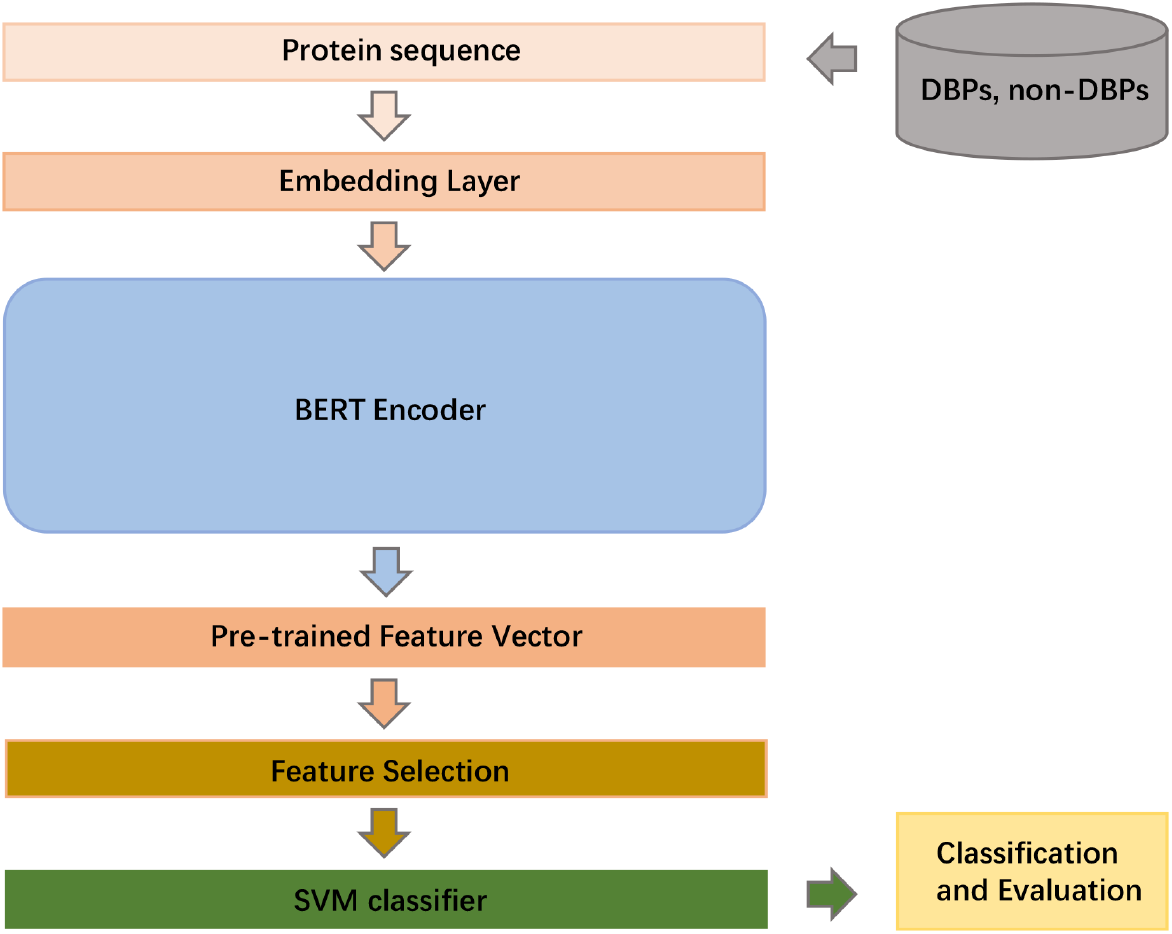
Pipeline of DBP2Vec. DBPs and non-DBPs are firstly fed into BERT Encoder to get the feature vectors, then optimal feature set are selected as input to SVM classifier for DBPs prediction.

ESM-1b is trained by using the masked language model BERT [46]. Each protein sequence is corrupted by replacing a fraction of the amino acids with a special mask token. The network is trained to predict the missing tokens from the corrupted sequence through a series of blocks that alternate self-attention with feed-forward connections. Self-attention allows the network to build up complex representations that incorporate context from across the sequence. Since self-attention explicitly constructs pairwise interactions between all positions in the sequence, the network directly represents residue–residue interactions in the protein sequence. Compared with popular PseAAC, ESM-1b captures more distant residue-residue interactions by using self-attention with fully connections. Moreover, ESM-1b learn biological properties (e.g., structural information) from large protein sequences database without any prior biological knowledge.

### Performance evaluation of DBP2Vec

To evaluate the performance of DBP2Vec, we use the standard performance metrics, such as sensitivity (SN), specificity (SP), accuracy (ACC), precision (PRE), F-score, AUC and MCC. These metrics can be calculated as follows:

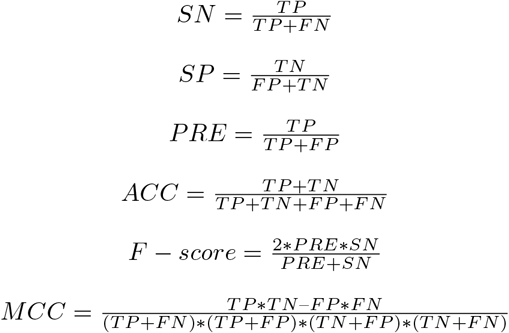

All the above metrics are based on the notions of TP, FP, TN, and FN, which correspond to number of positive samples identified correctly, negative samples identified incorrectly, negative samples identified correctly, and positive samples identified incorrectly, respectively. The MCC is an overall measurement of performance and another objective assessment index.

## Results

### Performance comparison on independent testing set PDB296

In order to verify the effectiveness of our proposed method, we compared our proposed method, DBP2Vec, with existing state-of-the-art methods, including TargetDBP [26], iDNAProt-ES [24], and DPP-PseAAC [19]. From Table 1, it is observed that DBP2Vec performs the best among the existing state-of-the-art methods on PDB269. The MCC of DBP2Vec are 0.631, an improvement of 0.097 over the second best result achieved by TargetDBP.

**Table 1.**
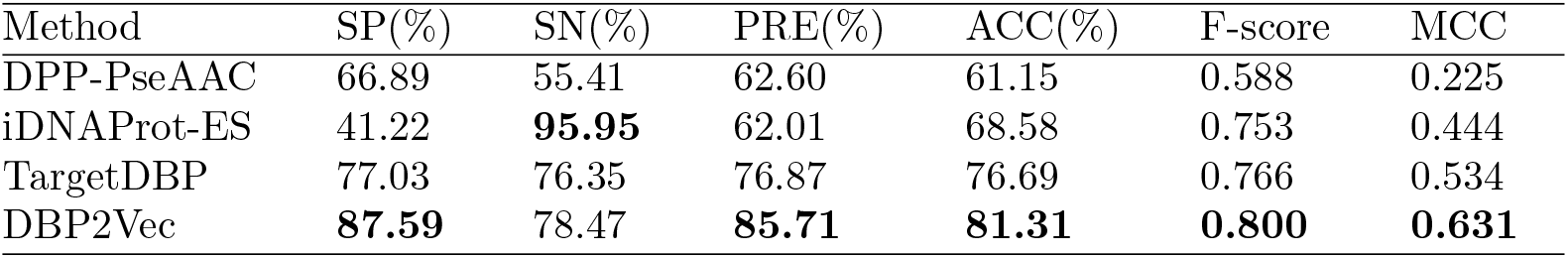
Comparison of DRBP2Vec and TargetDBP, iDNAProt-ES, DPP-PseAAC on PDB296

| Method | SP(%) | SN(%) | PRE(%) | ACC(%) | F-score | MCC |
| --- | --- | --- | --- | --- | --- | --- |
| DPP-PseAAC | 66.89 | 55.41 | 62.60 | 61.15 | 0.588 | 0.225 |
| iDNAProt-ES | 41.22 | **95.95** | 62.01 | 68.58 | 0.753 | 0.444 |
| TargetDBP | 77.03 | 76.35 | 76.87 | 76.69 | 0.766 | 0.534 |
| DBP2Vec | **87.59** | 78.47 | **85.71** | **81.31** | **0.800** | **0.631** |

### Performance comparison on independent testing set UniSwiss-Tst

In order to verify the effectiveness of our proposed method, we compared our proposed method, DBP2Vec, with existing state-of-the-art methods, including TargetDBP+ [27], iDNAProt-ES [24], and DPP-PseAAC [19]. From Table 2, it is observed that DBP2Vec performs the best among the existing state-of-the-art methods on UniSwiss-Tst. The MCC of DBP2Vec are 0.816, an improvement of 0.098 over the second best result achieved by TargetDBP+ [**?**].

**Table 2.** Comparison of DRBP2Vec and TargetDBP+, iDNAProt-ES, DPP-PseAAC on UniSwiss-Tst

| Method | SP(%) | SN(%) | PRE(%) | ACC(%) | F-score | MCC |
| --- | --- | --- | --- | --- | --- | --- |
| DPP-PseAAC | 55.64 | 54.59 | 55.32 | 55.25 | 0.550 | 0.105 |
| iDNAProt-ES | 80.84 | 73.75 | 79.38 | 77.30 | 0.741 | 0.486 |
| TargetDBP+ | 89.24 | 82.41 | 88.45 | 85.83 | 0.853 | 0.718 |
| DBP2Vec | **90.54** | **90.14** | **91.20** | **90.78** | **0.907** | **0.816** |

## CONCLUSION

In this paper, we proposed a novel DNA-binding proteins (DBPs) predictor (DBP2Vec) based on pre-trained protein language model (e.g., ESM-1b), which effectively exploit the context information of protein sequence. DBP2Vec is not only simpler, but also more accurate than existing state-of-the-art methods when tested on two benchmark DBPs datasets, e.g., PDB296 and UniSwiss-Tst, which verifies the effectiveness of pre-trained protein language model for DBPs prediction.

## Acknowledgments

We thank just about everybody.

## Notes

### Competing Interest Statement

The authors have declared no competing interest.

